# Modeling latent flows on single-cell data using the Hodge decomposition

**DOI:** 10.1101/592089

**Authors:** Kazumitsu Maehara, Yasuyuki Ohkawa

## Abstract

Single-cell analysis is a powerful technique used to identify a specific cell population of interest during differentiation, aging, or oncogenesis. Individual cells occupy a particular transient state in the cell cycle, circadian rhythm, or during cell death. An appealing concept of pseudo-time trajectory analysis of single-cell RNA sequencing data was proposed in the software Monocle, and several methods of trajectory analysis have since been published to date. These aim to infer the ordering of cells and enable the tracing of gene expression profile trajectories in cell differentiation and reprogramming. However, the methods are restricted in terms of time structure because of the pre-specified structure of trajectories (linear, branched, tree or cyclic) which contrasts with the mixed state of single cells.

Here, we propose a technique to extract underlying flows in single-cell data based on the Hodge decomposition (HD). HD is a theorem of vector fields on a manifold which guarantees that any given flow can decompose into three types of orthogonal component: gradient-flow (acyclic), curl-, and harmonic-flow (cyclic). HD is generalized on a simplicial complex (graph) and the discretized HD has only a weak assumption that the graph is directed. Therefore, in principle, HD can extract flows from any mixture of tree and cyclic time flows of observed cells. The decomposed flows provide intuitive interpretations about complex flow because of their linearity and orthogonality. Thus, each extracted flow can be focused on separately with no need to consider crosstalk.

We developed ddhodge software, which aims to model the underlying flow structure that implies unobserved time or causal relations in the hodge-podge collection of data points. We demonstrated that the mathematical framework of HD is suitable to reconstruct a sparse graph representation of diffusion process as a candidate model of differentiation while preserving the divergence of the original fully-connected graph. The preserved divergence can be used as an indicator of the source and sink cells in the observed population. A sparse graph representation of the diffusion process transforms data analysis of the non-linear structure embedded in the high-dimensional space of single-cell data into inspection of the visible flow using graph algorithms. Hence, ddhodge is a suitable toolkit to visualize, inspect, and subsequently interpret large data sets including, but not limited to, high-throughput measurements of biological data.

The beta version of ddhodge R package is available at: https://github.com/kazumits/ddhodge

## 1. Methods

### 1.1 Hodge decomposition on graph

Our ddhodge is built on a mathematical framework of the Hodge decomposition (HD) that provides an analogy of gradient, curl, and harmonic (cyclic) flows of the vector field. Jiang et. al proposed HodgeRank [1] as an application of HD to reconstruct global rankings from the pairwise relations of user ratings on Netflix movies. We developed the R package ddhodge which implemented the HD framework with the coboundary operators on a graph (grad, div, curl, and Laplacians) as introduced in HodgeRank and the introductory notes of Hodge Laplacians on graphs [2]. We briefly explain the results of HD and some notations used in this article.

The cochains *C*^0^, *C*^1^, and *C*^2^ are the spaces of functions that assign a real value to the cartesian product of vertices in *V*; *V* → *ℝ*, *V* × *V* → ℝ and *V* × *V* × *V* → ℝ, respectively. The coboundary operators *δ*_0_ = grad and *δ*_1_ = curl are the maps between the cochains *C^k^* → *C*^k+1^, and their adjoints *C^k^* → *C*^*k*-1^ are 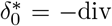 and 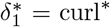 which are defined under an inner product 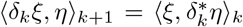 where *ξ* ∈ *C^k^* and *η* ∈ *C*^*k*+1^. They hold the relation 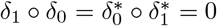. HD states that the orthogonal decomposition of *C^k^* exists as:

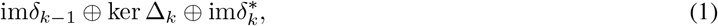

where 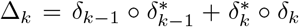 is Hodge Laplacian. If *k* = 1, the left term is the space of the gradient flow, and the central and right terms are those of divergence-free (div*X* = 0 iff *X* ∈ im(grad)^⊥^) harmonic and curl flow, respectively. If 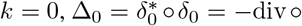 grad is known as graph Laplacian in graph theory, and *C*^0^ is decomposed into ker *δ*_0_ and 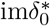. We illustrate the result of HD in Figure 1.

**Figure 1:**
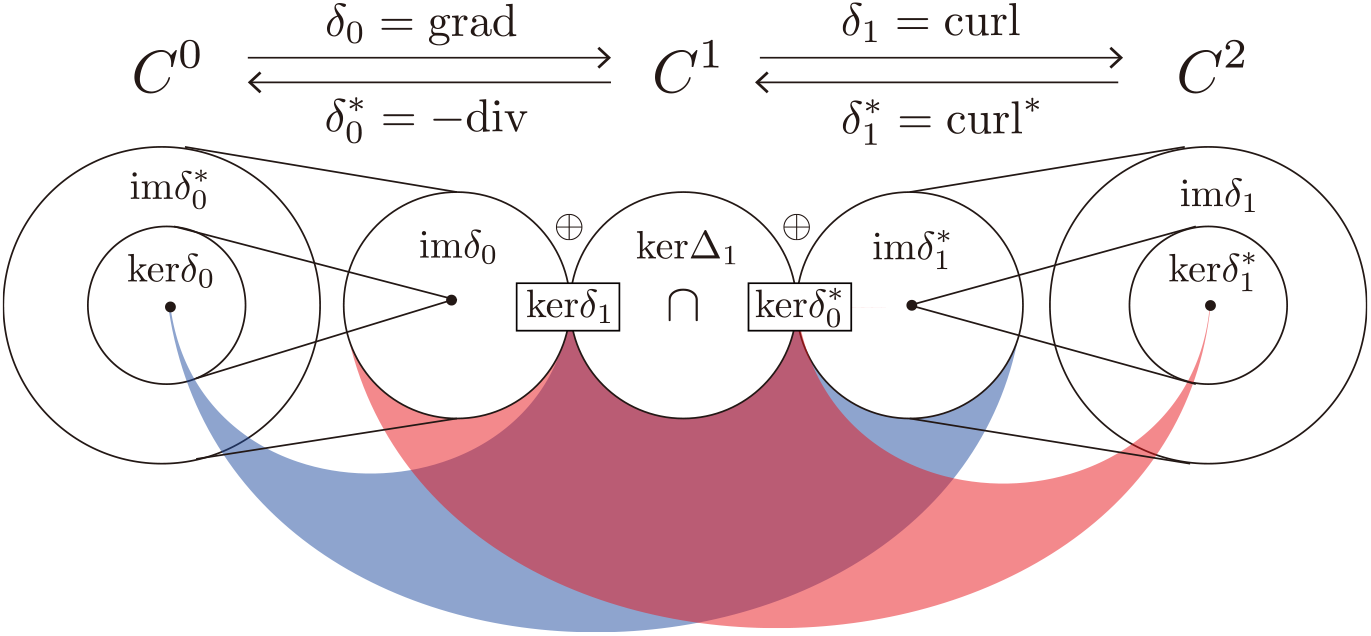
The scheme of Hodge decomposition.

The cochains *C*^0^, *C*^1^, and *C*^2^ can be viewed as vector spaces of weighted nodes, edges, and triangles in graph *G*. Accordingly, the operators of grad, div, and curl are represented as matrices (linear maps between the weights of nodes, edges, and triangles; see [1, 2]. Using the HD setup on the graph, some flow decomposition problems of *G* are formulated as a simple least square problem, such as min_*b*_ ||*y* – grad*b*||^2^ where *y* is the given flow (weighted edges). A solution *b* is obtained by the orthogonal projection of *y* onto the image of grad, and is interpreted as a potential field that produces gradient flows in *y*. Then, the residual, consisting of the curl and harmonic flow, is divergence-free as explained above.

### 1.2 Designing edges through diffusion maps

The setup of HD was completed as above, but the problem of what kind of edges (direction and weights) would be appropriate for cells still remained. The Waddington epigenetic landscape [3] is a famous picture of cell lineage specification and has interested researchers who explore the kinetics that determine cell fates [4, 5]. This assumes that a stochastic process along the energy potential of a biochemical system is a natural way to model cell fate. We employed the diffusion process as a candidate of underlying dynamics that produces the time structure in single-cell (sc)RNA-seq data because it is a simple (requires few assumptions) and straightforward way to model the development of the state of cells (e.g., the gene expression profile) as entropy increases.

Diffusion maps [6, 7] are a suitable method of nonlinear dimension reduction that accounts for random walk on data points and was also applied for scRNA-seq data [8, 9]. In diffusion maps, a low-dimensional representation of the data is obtained through the spectral decomposition of transition probability matrix *M. M* is obtained by normalizing the affinity matrix (each row sums to 1), which is constructed from a defined similarity of data points. We used Gaussian affinity exp(||*x_i_* – *x_j_*||^2^/*σ_i_σ_j_*) with the local scaling parameter *σ_i_* that is determined by the distance to the *n*-th nearest neighbor from *x_i_* [10]. Local scaling equalizes the different scales of within-cluster variations, which are seen in the mix of slow and fast transitions of states.

The vector field which corresponds to the diffusion process at a fixed time *t* = *s* is a potential field that contains only gradient flow. We designed the edges as the gradient arose from a given potential (concentration) at time *t* = *s*. The initial condition at *t* = 0 (concentrations of starting cells) is given by any prior data knowledge, such as the identification of a cell population with marker genes, known cell groups before differentiation stimuli occur, or cell groups determined by clustering methods if an unbiased manner is preferable. The potential *u* at *t* = *s* is calculated as:

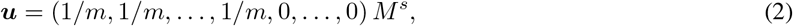

where the first *m* elements are selected as the starting cell. The edges are defined by gradu (i.e., the differences: *u_i_* – *u_j_*). We refer to this as the diffusion graph. The directed acyclic graph (DAG) can be constructed as above, though visualization and inspection of the graph to understand the diffusion dynamics remain difficult because it is a fully-connected complete graph (*N*^2^ edges for *N* cells). We therefore keep only the edges of *k*-nearest neighbors (NN) on the diffusion coordinate. *k*-NN graph construction is frequently used to obtain sparse representations of graphs [11, 12]. However, pruning by *k*-NN criteria breaks several good properties in the original diffusion graph such as divergence and potential. We therefore appended the additional step after edge pruning to keep these properties while maintaining the sparsity of the graph as explained below.

## 2. Recovering the diffusion nature using a sparse network

The HD framework is intuitive, so helps us to consider several extensions such as placing constraints on the qualitative characteristics of graph *G*. To preserve the original diffusion, we therefore chose to keep divergence because this tells us the source and sink information of a flow. We constructed the objective function to minimize the loss of original divergence using the pruned graph as:

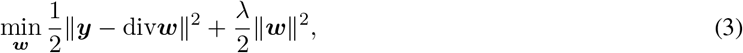

where *y* is the divergence of the original diffusion graph, and div is the divergence operator of the pruned *k*-NN graph. Finally, we obtained the edges of the sparse graph consisting of pure gradient flow (DAG) by the orthogonal projection of the solution *w* onto im(grad) by:

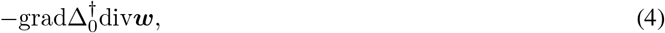

where 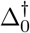 is the Moore–Penrose pseudoinverse of the graph Laplacian. The residual parts of *w* consisting of curl- and harmonic-flow can be safely ignored because they do not contribute to divergence. We also note that once *w* is fixed, the recovered potential 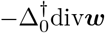 is not guaranteed to be close to the original one simply because the graph structure (Δ_0_) is different in the *k*-NN graph.

Instead of the regression as above, the generalized lasso [13] formulation can be incorporated to directly solve the problem of the sparse construction of gradient flows. However, the computational cost may be infeasible.

### 2.1 10 steps for sparse diffusion graph construction

We summarize our construction of the sparse diffusion graph in 10 steps as follows:

1. Normalize the count matrix using medians of ratio normalization [14]
2. Perform variance-stabilizing transformation using log_2_ (normalized count + 0.5)
3. Reduce to p dimensions using principal component analysis
4. Construct a transition probability matrix using Gaussian affinity with local scaling
5. Select starting cells and proceed the diffusion for *s* step (eq:2)
6. Construct a fully-connected diffusion graph equipped with gradient flow as edges
7. Calculate the divergence of the original diffusion graph
8. Prune the network using *k*-NN criteria at specified *k* on diffusion coordinates
9. Recover the original divergence using ridge regression (eq:3)
10. Construct a sparse diffusion graph using reconstructed gradient flow (eq:4)

**Figure 2:**
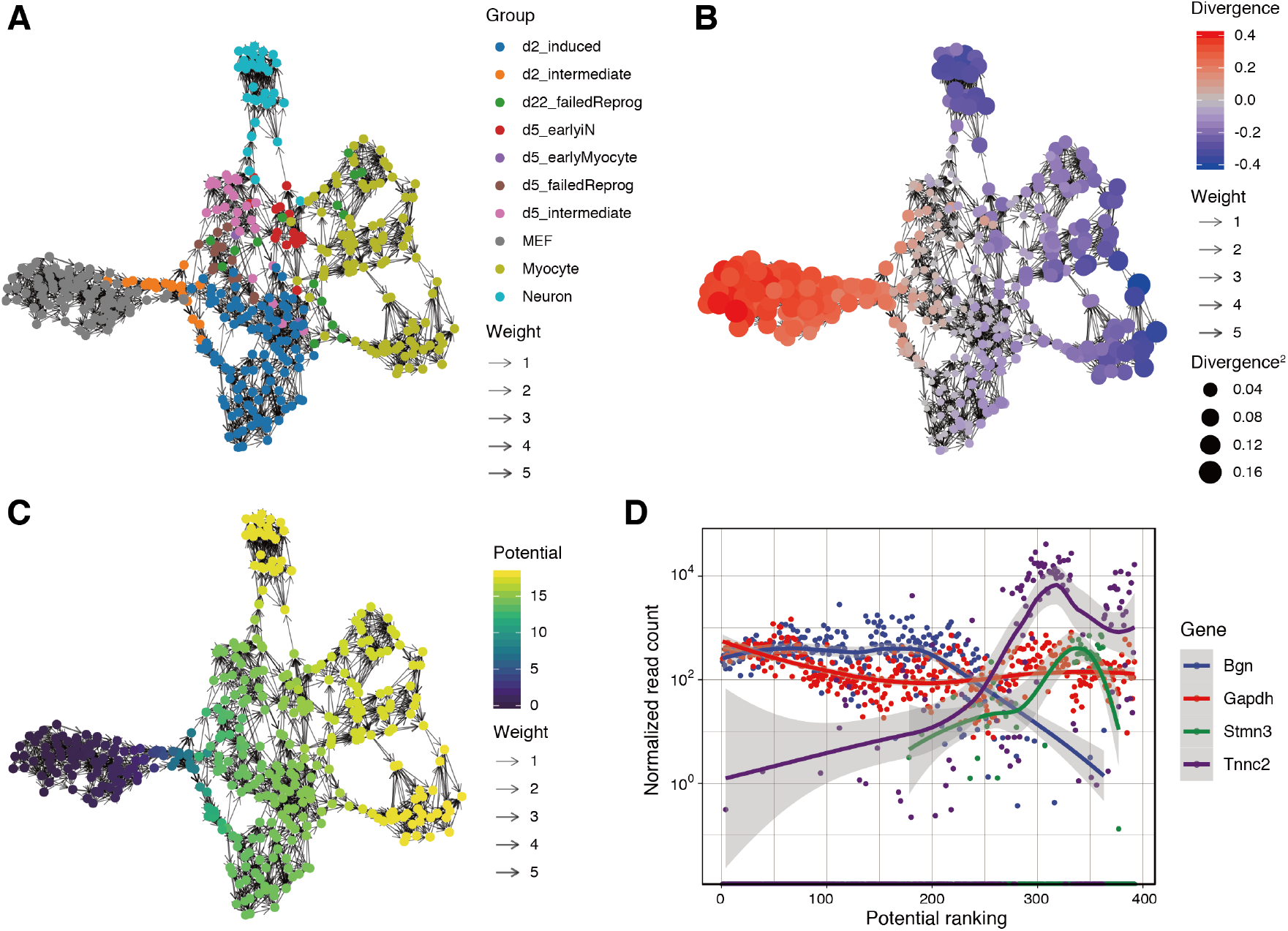
Visualization with group labels (A), divergence (B), potential (C), and potential-ordered representative gene expression (D) with smoothed curves. The force-directed layout was used.

## 3. Results

We first present the visualization performance of ddhodge using scRNA-seq in [15]. The data are a collection of single-cell gene expression profiles collected at several time points in the direct reprogramming of mouse embryonic fibroblasts (MEF). Figure 2A shows visualization by the divergence-recovered sparse diffusion graph. Our ddhodge with diffusion successfully clustered the cell lineages without using the information of group labels (colored circles).

We also visualized the recovered divergence in the same layout (Figure 2B). The divergence informs us that the “source” cells that we provided as start cells are in red (divergence > 0) as expected. The negative divergence (blue circles) indicated the existence of corresponding “sink” cell populations at the terminal points of the path to the myocyte and neuron groups from MEFs. The gray cells (around zero divergence) can be reasonably interpreted as waypoints. The potential, which corresponds to the proportion of cells in each state at time *t* = *s*, can be used as “pseudo-time” information (Figure 2C, D). A human-friendly tree layout is accessible because the graph is purely DAG (Figure 3A).

As described above, it is straightforward to follow the path of reprogramming along the arrows in the sparse diffusion graph. The visualizations show multiple paths from MEFs to neurons and myocytes. The maximum flow of the graph can be used to extract a path that maximizes the flow between specified node pairs with given capacities of edges (gradients) (Figure 3B). The flow can be seen as a representative path in the reprogramming.

## 4. Discussion

We developed ddhodge as a framework to dissect the latent flows embedded in a collection of single cells. Our analysis of non-linear trajectories of high-dimensional single-cell data was transformed into the traversing of sparse diffusion graphs. The graphs enable the unveiling of directions in differentiation, which are visualized as simplified flows. These are useful to identify the path to a specific differentiation or the appearance of a particular cell population such as the stem cell state, intermediate state, or terminally-differentiated state.

**Figure 3:**
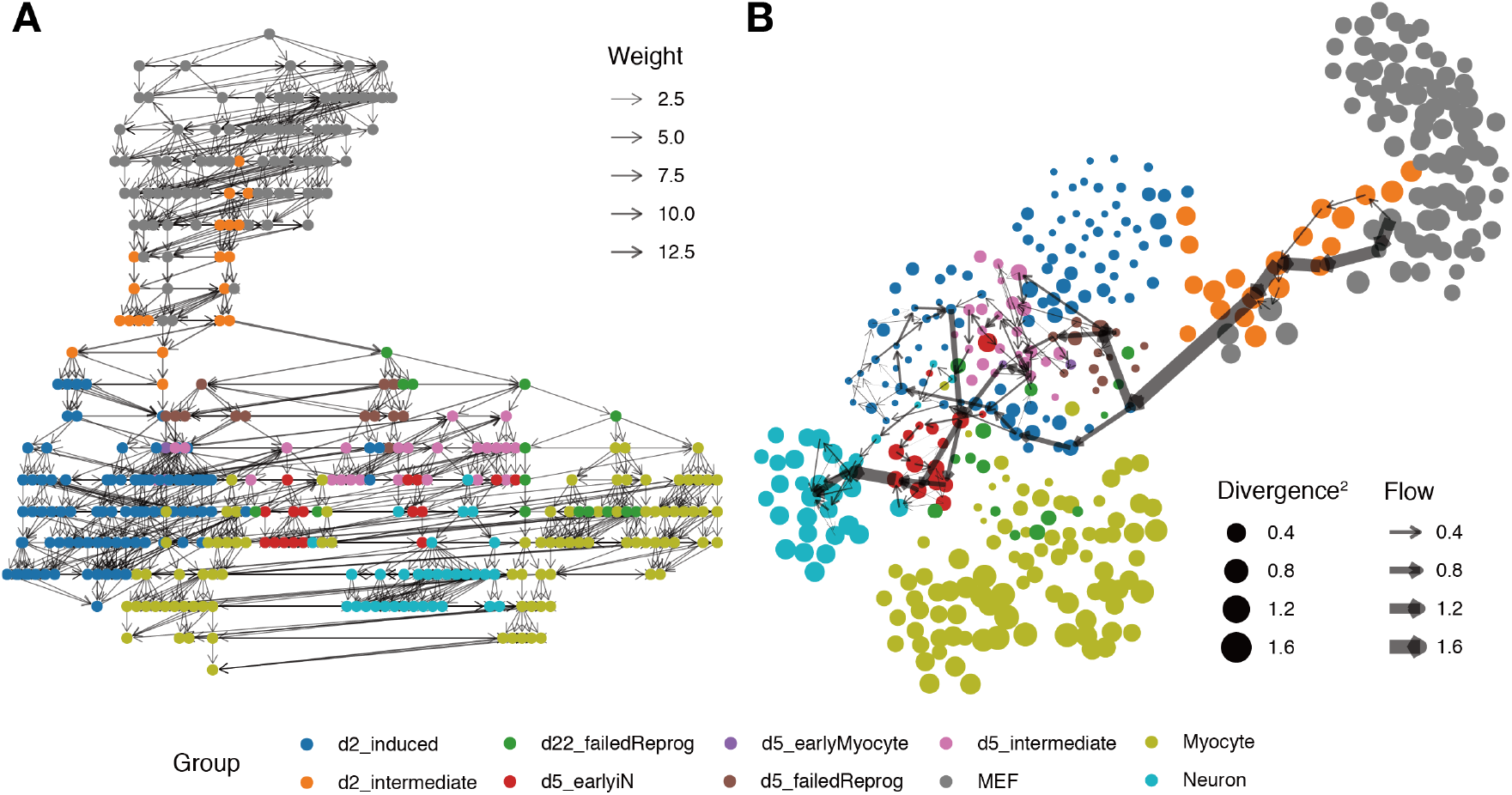
Tree layout (A) and max-flow between a specified source and sink pair (B; large graph layout).

Monocle [16] has become a widely-used tool for scRNA-seq to trace lineage-specific differentiation. Several alternative approaches have been proposed for the trajectory analysis [11, 8, 17, 12], and these are comprehensively reviewed in [18, 19, 20]. Current approaches for trajectory analysis assume that the data contain either branched (generally a tree) or cyclic structures [21, 22, 23]. However, the only requirement for HD is that the graph is directed; it also has the unique advantage of the flexibility of handling any vector field.

The development of HD-based single-cell trajectory analysis is still in its infancy, although it has succeeded in extracting the flows in differentiation paths from scRNA-seq data. However, understanding the more realistic nature of temporal regulation of cells, including circadian rhythms and cell cycles, is still challenging because the information of space and time is not contained in single-cell profiles. Although the methodology uses the similarity of measured data points, this does not deduce causal directions among cells. Designing edges using emerging technologies of statistical causal inference [24, 25, 26, 27] combined with single-cell omics data other than transcriptomes, e.g., epigenomes [28] could pave the way for the reconstruction of causal flows including cyclic relations.

